# Heterogeneity of brain dynamics in genetic and psychiatric conditions

**DOI:** 10.64898/2026.09.01.748394

**Authors:** Adrien E.E. Dubois, Sarah Lippé, Charles-Olivier Martin, Inga S. Knoth, Valérie K. Fontaine, Anne-Marie Bélanger, Pascale Abadie, Melissa T. Carter, Mélanie Couture, Mayada Elsabbagh, Alan Evans, Carl Ernst, Baudouin Forgeot d’Arc, Ridha Joober, Guy Rouleau, Julie Scorah, Christine Lucas Tardif, Ma’n H. Zawati, Grace C. Westerkamp, Ernest V. Pedapati, Craig A. Erickson, Borja Rodriguez-Herreros, Nadia Chabane, Kenza Latrèche, Marie Schaer, Aline Lefebvre, Richard Delorme, Sara Jane Webb, James C. McPartland, Sébastien Jacquemont, Guillaume Dumas

## Abstract

Whether the heterogeneity of psychiatric conditions converges on shared neurophysiological alterations or translates into distinct signatures remains unclear. We assembled high-density electroencephalogram (hd-EEG) resting-state recordings from 4,812 individuals aged 5 months to 66 years across 11 psychiatric conditions, a broad spectrum of rare genetic variants, and typically developing (TD) individuals. We established normative developmental trajectories of source-space EEG across spectral organization, connectivity, and signal complexity. Psychiatric conditions showed small deviations, revealing a shared transdiagnostic profile. In contrast, single rare variants showed substantially larger, distinct and sometimes mirror-opposite signatures that collapsed toward the psychiatric profile when pooled. Autism Spectrum Disorder showed some of the smallest group-level effects yet the largest individual deviations, indicating substantial but directionally inconsistent alterations. EEG deviations followed a cortical gradient, with larger effects in sensorimotor regions. We demonstrate that sample sizes in the hundreds are required for robust associations with psychiatric diagnoses. This interactive open resource provides normative scores to benchmark future results.

## Introduction

Psychiatric conditions are heritable disorders arising from disruptions to brain development and collectively represent a major global health priority^1–3^. Despite substantial advances in genetics, neuroscience, and clinical characterization, no robust neurophysiological biomarker has been established for any major condition, including autism spectrum disorder (ASD)^4^. Current diagnosis therefore remains dependent on behavioral assessment alone. One major obstacle to biomarker discovery is the profound heterogeneity, which spans diverse behavioral phenotypes, developmental trajectories, and genetic architectures while exhibiting extensive overlap across diagnostic categories^5,6^.

Electroencephalography (EEG) provides a promising avenue for addressing this challenge. Resting-state EEG captures neural dynamics with millisecond temporal resolution, is inexpensive and scalable, and can be acquired across the lifespan with minimal task demands, making it particularly well suited for large transdiagnostic studies. Neural oscillations are thought to support information processing and cortical communication through the synchronization of distributed neural activity^7,8^. Prior work has reported alterations across multiple electrophysiological domains in psychiatric conditions, including oscillatory power^9^, functional connectivity^10,11^, and signal complexity ^12^. However, findings have often been inconsistent across studies, likely reflecting methodological heterogeneity, limited sample sizes, and the biological diversity encompassed within current diagnostic categories^13,14^. As a result, it remains unclear whether EEG alterations reflect condition-specific signatures or shared disruptions of neural dynamics across conditions.

The challenges associated with heterogeneity may partly arise from the mismatch between behavioral diagnoses and underlying biology. Psychiatric conditions such as ASD and attention-deficit/hyperactivity disorder (ADHD) aggregate individuals with highly heterogeneous genetic etiologies, whereas individuals with the same rare high-impact genetic variants define more biologically homogeneous groups^15,16^. EEG measures are themselves strongly heritable, particularly in the alpha band, with twin studies attributing a large proportion of variance to additive genetic influences^17,18^. Yet, the majority of EEG studies in psychiatric conditions have focused on broad clinical diagnoses, while only a few have focused on specific genetic syndromes, preventing direct comparison between psychiatric diagnostic categories and associated genetic predispositions. A few rare copy number variants (CNVs) and monogenic syndromes such as Fragile X syndrome (FXS) have been associated with marked alterations in spectral power and functional connectivity^19–26^, yet these studies focused on very few genomic loci and genes (i.e., 16p11.2, 22q11.2, 15q11-13, and FMR1) and the findings are fragmented across studies and features. Whether neurophysiological alterations converge or diverge across psychiatric disorders and genetic etiologies remains unresolved.

Here, we assembled the largest transdiagnostic and cross genetic-predisposition resting-state EEG dataset to date, comprising 4,812 high-density recordings from individuals spanning 11 psychiatric conditions and rare genetic variants across the lifespan. Genetic groups range from homogeneous (individuals carry the same genetic predisposition) to increasing levels of genetic heterogeneity (different classes of variants disrupting different neurodevelopmental genes).

The overarching aim is to establish lifespan charts to investigate EEG alterations/deviations across psychiatric conditions and related genetic predisposition, as well as the impact of heterogeneity on our ability to identify associations.

Using normative source-space modeling^27,28,28–31^, we characterized developmental trajectories of spectral, connectivity, and entropy features in a typically developing (TD) reference population and quantified individual-level deviations across clinical and genetic groups. We then tested whether electrophysiological alterations converge across behaviorally defined diagnoses, whether genetically homogeneous groups exhibit more distinct signatures, and whether these deviations share a common cortical organization relative to established cortical gene expression gradients. Finally, we evaluated the reproducibility of EEG effect sizes across sample sizes to establish empirical benchmarks for biomarker discovery in neurodevelopmental research.

## Results

### Source-reconstructed EEG features across a transdiagnostic cohort

The combined sample comprised 4,812 participants drawn from 9 datasets spanning an age range of 5 months to 66 years, all acquired using the same high-density 128-channel EGI system (Figure 1A). Comorbid diagnoses most frequently involved ADHD and ASD (Figure 1B). Signal quality, indexed by the proportion of bad segments, differed significantly across conditions and from TD (Figure 1C) and was included as a covariate in subsequent analyses.

**Figure 1.**
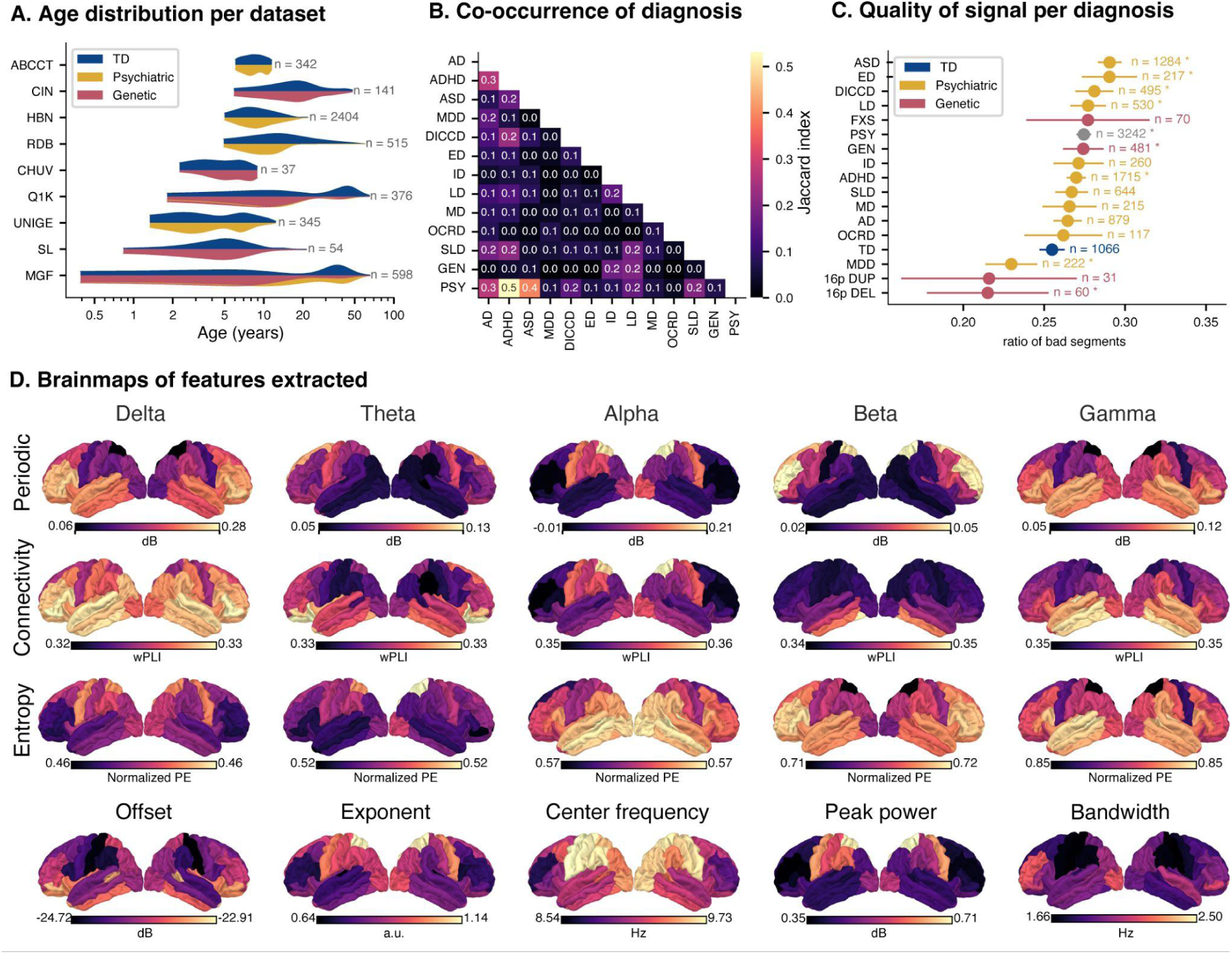
Sample characteristics and spatial distributions of EEG source features. **A**. Age distribution across recording sites. Each row shows the density for one database, sorted by median age. Highlights heterogeneity in developmental coverage across datasets. **B.** Pairwise Jaccard index between diagnostic categories. Each cell represents the proportion of subjects carrying both diagnoses, relative to those carrying either diagnosis. Values range from 0 (no overlap) to 1 (identical groups). **C.** Ratio of bad segments per diagnostic group (± 95% CI), sorted by recording quality. Identifies whether specific populations are systematically harder to record, informing potential quality-related confounds in downstream analyses. **D.** Source-level spectral, connectivity, and entropy measures in the typically developing (TD) group. Spectral features were decomposed into aperiodic components (offset and exponent), alpha-band components (center frequency, peak power, and broadband power), and periodic power. Periodic power, entropy, and connectivity were analyzed within canonical frequency bands: delta (2–4 Hz), theta (4–8 Hz), alpha (8–12 Hz), beta (12–30 Hz), and gamma (30–45 Hz). Abbreviations: ABCCT: Autism Biomarkers Consortium for Clinical Trials, AD: Anxiety Disorder, ADHD: Attention-Deficit/Hyperactivity Disorder, ASD: Autism Spectrum Disorder, CHUV: Centre Hospitalier Universitaire Vaudois, CIN: Cincinnati Children’s Hospital, DEL: deletion, DICCD: Disruptive, Impulse-Control, and Conduct Disorders, DUP: duplication, ED: Elimination Disorders, FXS: Fragile X Syndrome, GEN: genetic, HBN: Healthy Brain Network, ID: Intellectual Disability, LD: Language Disorder, MD: Motor Disorders, MDD: Major Depressive Disorder, MGF: Montreal Genetic Family, OCRD: Obsessive-Compulsive and Related Disorders, PSY: psychiatric conditions, Q1K: Quebec 1000, RDB: Robert Debré University Hospital Center, SL: Simons Searchlight, SLD: Specific Learning Disorder, TD: Typically Developing, UNIGE: Université de Genève, 16p DEL: 16p11.2 deletion, 16p DUP: 16p11.2 duplication.

Source reconstruction was performed for three classes of features: the power spectrum decomposed into aperiodic and periodic components (see Figure S5 for the power spectrum and its fit), functional connectivity, and signal entropy. As expected, the spatial distributions of periodic signals showed the characteristic posterior alpha rhythm and frontal dominance in the delta / gamma bands (Figure 1D) and the aperiodic exponents showed parietal localization, while offsets showed frontal predominance. The median alpha center frequency was 9.39 Hz (peak power: 0.50 dB; bandwidth: 1.83 Hz), consistent with normative values for this age range.

Source reconstruction was validated against independent electrophysiological data by correlating spatial patterns of periodic power with the electrophysiological neuromaps^32,33^. Significant positive correlations were observed across frequency bands (delta: r = 0.66*; theta: r = 0.32*; alpha: r = 0.65*; beta: r = 0.41*), supporting the biological validity of the reconstructed oscillatory maps. Gamma-band activity showed a stronger correspondence with high-gamma (r = 0.59*) than with low-gamma (r = 0.17) electrophysiological patterns.

Sensitivity analysis showed that alternative methods to decompose periodic and aperiodic signals did not influence these results (Figure S1). Separating the alpha and beta bands into low- and high-frequency components revealed distinct spatial organizations (see Figure S2); however, to maintain comparability with prior studies, analyses were performed using the five canonical frequency bands.

Spatial distribution of mean connectivity and entropy have not been previously characterized. It was largely similar across frequency bands, except for higher values in the alpha band. Alpha-band connectivity was elevated in posterior parietal and occipital regions relative to frontal regions, consistent with posterior alpha dominance^34^. Entropy showed a strong frequency-dependent hierarchy, increasing systematically from delta to gamma, reflecting frequency-specific signal complexity. Spatial differences displayed distinct topographies, with delta and theta exhibiting parietal dominance and a progressive shift toward fronto-temporal dominance from alpha to gamma.

### Source level EEG lifespan charts reveal feature-specific age-related changes

We characterized EEG lifespan variations for the 20 features using GAMLSS (Generalized Additive Models for Location, Scale and Shape)^35^.

The most pronounced changes occurred between 0 and 20 years of age. Periodic power and connectivity across the full frequency range (2–45 Hz) decreased with age, in contrast to entropy (Figure 2A). Individual frequency bands revealed distinct developmental trajectories (Figure 2B). We observed the expected age-related reduction in periodic power in the delta and theta bands, along with an increase in the alpha band. The latter was accompanied by a reduction of alpha peak and bandwidth as well as a shift towards higher frequencies. Age-related effects of beta and gamma power, previously less documented, demonstrated respectively an increase and a decrease with age.

**Figure 2.**
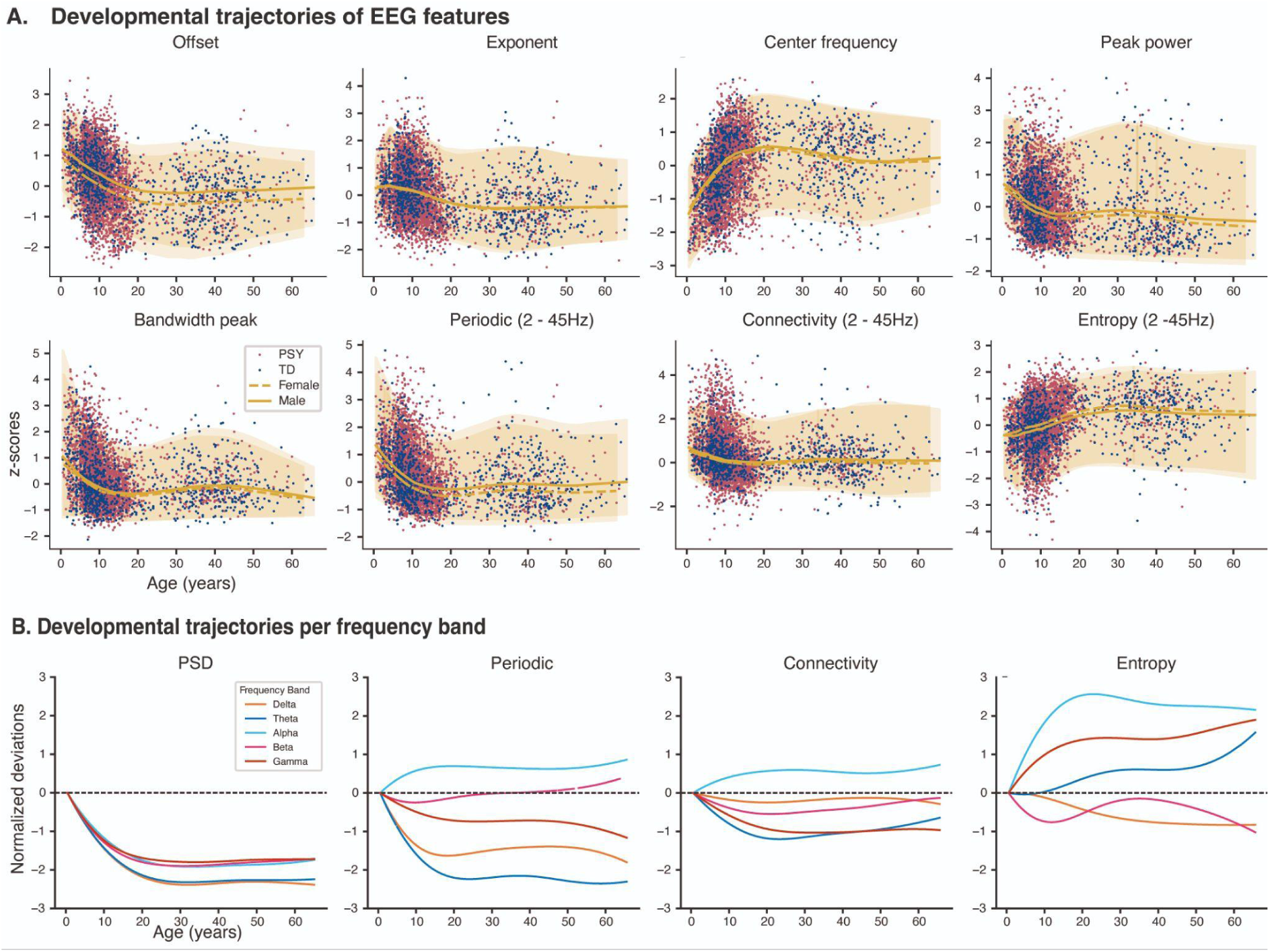
EEG developmental trajectories. **A.** Normative developmental trajectories in typically developing (TD) participants for all extracted EEG features. Spectral features were decomposed into aperiodic components (offset and exponent), alpha-band components (center frequency, peak power, and broadband power), and periodic power. Periodic power, connectivity, and entropy were computed across the full frequency spectrum (2–45 Hz). **B.** Normative developmental trajectories in TD participants, expressed in standard units and aligned to start at 0 at the youngest age to prevent overlapping across frequency bands. Power spectral density (PSD), periodic power, entropy, and connectivity were analyzed within canonical frequency bands: delta (2–4 Hz), theta (4–8 Hz), alpha (8–12 Hz), beta (12–30 Hz), and gamma (30–45 Hz). All lifespan trajectories at the ROI level, along with the corresponding model quality metrics, are available at https://dub21.github.io/D-EEG/

Aperiodic components decreased substantially, reflecting a broad reduction in overall spectral power and a relative increase in higher-frequency compared with slower-frequency activity across development. Offset was the feature with the largest sex-related effects (increased in males).

Of note, the power spectral density (PSD) age trajectories were primarily driven by the offset, highlighting the added value of decomposition. Connectivity showed the least age-related effects. Entropy showed a distinct developmental profile from spectral power and connectivity, with marked age-related increases in entropy in the alpha band, followed by gamma and theta, while delta and beta showed decreases.

### EEG signal effect size and heterogeneity across psychiatric conditions and specific variants

Based on the normative trajectories of the 20 features, we analyzed deviations across all conditions (Figure 3A). For psychiatric conditions, the largest proportion of significant and convergent associations was observed for EEG spectral features, particularly periodic power (Figure S3). These deviations followed a broad U-shaped pattern characterized by positive deviations in the delta and gamma bands and reduced values in the alpha and beta bands. Similarly, a shift in the alpha peak toward lower frequencies and a broader bandwidth were observed across all psychiatric conditions, along with reduced aperiodic components. Entropy displayed a comparable U-shaped pattern across diagnoses, with the trough shifted toward the theta and alpha bands. Connectivity showed the smallest proportion of significant associations and convergence, although theta-band connectivity was consistently increased across conditions.

**Figure 3.**
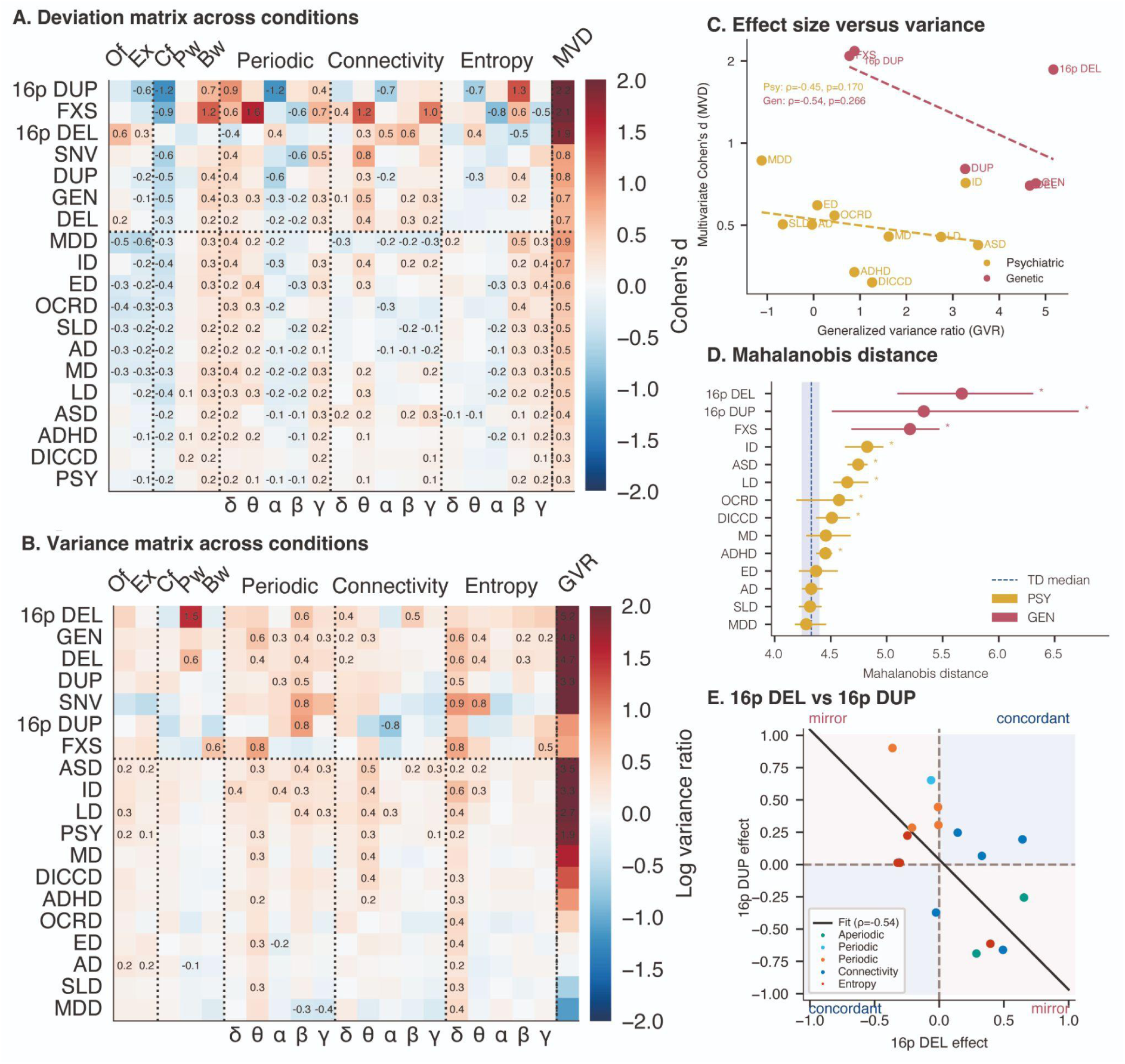
Multivariate EEG Deviations Across Conditions. **A.** Each cell shows Cohen’s d between cases and TD for the corresponding EEG feature group. Positive and negative values indicate the direction of the difference relative to TD. Values are displayed only when FDR-significant (q < 0.05), as assessed by permutation testing. Conditions are ranked within genetic and psychiatric categories by the multivariate Cohen’s d (MVD, rightmost column), which quantifies the shift in group centroid across all EEG features simultaneously. **B.** Each cell shows the log variance ratio between conditions and TD for the corresponding EEG feature. Positive and negative values indicate greater and lower inter-subject variability, respectively, in the clinical group relative to TD (a value of 0 corresponds to control-level variance). Conditions are ranked by the generalized variance ratio (GVR, rightmost column), which quantifies the difference in overall multivariate dispersion between groups as a bias-corrected log ratio of covariance matrix pseudo-determinants. Note that GVR and individual-feature log variance ratios share the same log-ratio scale but are not directly numerically comparable, as GVR integrates dispersion across all features simultaneously. Values are displayed only when FDR-significant (q < 0.05), as assessed by Levene’s test for individual features and by permutation testing for GVR. **C.** Each point represents a condition. The x-axis shows the generalized variance ratio (GVR), a bias-corrected log ratio of multivariate dispersion relative to TD (GVR = 0 indicates control-level dispersion). The y-axis shows the multivariate Cohen’s d (MVD) on a logarithmic scale, quantifying the shift in group centroid relative to TD. Psychiatric conditions are shown in yellow and genetic conditions in red. Dashed lines show separate Spearman regression fits for psychiatric and genetic conditions. **D.** *Each dot represents the median Mahalanobis distance computed in the TD reference space, with horizontal bars indicating the 95% bootstrap confidence interval (n = 2,000 resamples). The dashed line and shaded band indicate the median and 95% bootstrap CI of the TD reference group, respectively. Psychiatric and genetic conditions are shown in distinct colors and sorted by ascending median distance. : significant difference from TD after FDR correction (α = 0.05; permutation test). **E.** The scatterplot compares Cohen’s d for the 16p DUP group (y-axis) and the 16p DEL group (x-axis) across EEG features. Background shading indicates the direction of deviations relative to TD: red indicates opposite-direction deviations, whereas blue indicates concordant deviations. The black line shows the linear regression fit across all EEG features; Pearson’s r is reported in the legend. Abbreviations: AD: Anxiety Disorder, ADHD: Attention-Deficit/Hyperactivity Disorder, α: alpha, ASD: Autism Spectrum Disorder, β: beta, Bw: bandwidth, Cf: center frequency, DEL: deletion, δ: delta, DICCD: Disruptive, Impulse-Control, and Conduct Disorders, DUP: duplication, ED: Elimination Disorders, Ex: exponent, FXS: Fragile X Syndrome, γ: gamma, GEN: genetic, GVR: Generalized Variance Ratio, ID: Intellectual Disability, LD: Language Disorder, MD: Motor Disorders, MDD: Major Depressive Disorder, MVD: Multivariate Cohen’s d, OCRD: Obsessive-Compulsive and Related Disorders, Of: offset, PSY: psychiatric conditions, Pw: peak power, SLD: Specific Learning Disorder, TD: Typically Developing, θ: theta, 16p DEL: 16p11.2 deletion, 16p DUP: 16p11.2 duplication.

The same periodic U-shaped pattern was also observed across genetic variant groups, with the exception of an inverted U-shaped pattern in 16p deletion carriers. Significance and convergence were reduced for the aperiodic components and entropy. Conversely, connectivity showed greater significance and convergence in genetic variant groups than in psychiatric conditions. Considering all the features, we observed a correlation between psychiatric conditions and genetic groups (Figure S7) while some specific variants such as the 16p deletion and duplication manifested opposite patterns (Figure 3E).

We hypothesized that heterogeneity across psychiatric conditions and genetic variant groups would be negatively correlated with overall EEG effect sizes. Consistent with this hypothesis, effect sizes for the 20 univariate EEG features, expressed as Cohen’s *d*, were two- to four-fold smaller in psychiatric conditions (maximum *d* < 0.5) than in groups of individuals carrying the same genetic variant (up to *d* = 1.5) (Figure 3A). The same pattern was observed for multivariate Cohen’s *d*, which revealed substantially larger deviations than the univariate measures. Conditions displayed differences in variance, from low variance in MDD to high variance in ASD. Similarly, genetic variants range from small variance for FXS and 16p11.2 duplication to high variance for 16p11.2 deletion (Figures 3B and 3C).

To further investigate the heterogeneity hypothesis for conditions with very low deviations (i.e., ASD), we computed a Mahalanobis distance across features. This allows to detect any deviation across features or individuals without the constraint of a homogeneous group effect. Results demonstrate that ASD showed the largest and most significant (with ID) Mahalanobis distance from TD suggesting large and heterogeneous individual deviations (Figure 3D). This contrasts with MDD, which showed the largest Cohen’s d with the smallest Mahalanobis distance.

Sensitivity analysis demonstrates that data quality (Figure S4) or spectral fit parametrization (Figure S5) do not influence these results.

### EEG deviations align with the cortical gradient

We investigated whether EEG deviations exhibited a structured spatial organization across the cortex. To address this question, we used ROI-level normative models and examined their spatial correspondence with cortical gradients. Specifically, we tested the spatial similarity between deviation maps and the first principal component of cortical gene expression (PC1 gene expression), a well-established cortical gradient that orders regions from primary sensorimotor to higher-order association cortices32,36. Mean (Figure 4B) and variance (Figure 4A) maps of absolute effect sizes were computed across diagnoses to obtain consensus maps. When averaged across features, the mean deviation map showed a significant correlation (r = 0.58) with the cortical gene expression gradient (Figure 4B), largely driven by periodic activity (r = 0.7, Figure 4C). Regions exhibiting the largest deviations were concentrated in sensorimotor cortices, whereas regions with greater variance tended to be located in association cortices; however, the corresponding variance effects were small and did not survive spin permutation testing. The association between deviation maps and the cortical gene expression gradient was preserved across diagnoses (Figure 4D) and ASD datasets (Figure 4E). When using the knee parameterization, part of the spatial alignment with the cortical expression gradient was accounted for by the knee component (See Figure S6).

**Figure 4.**
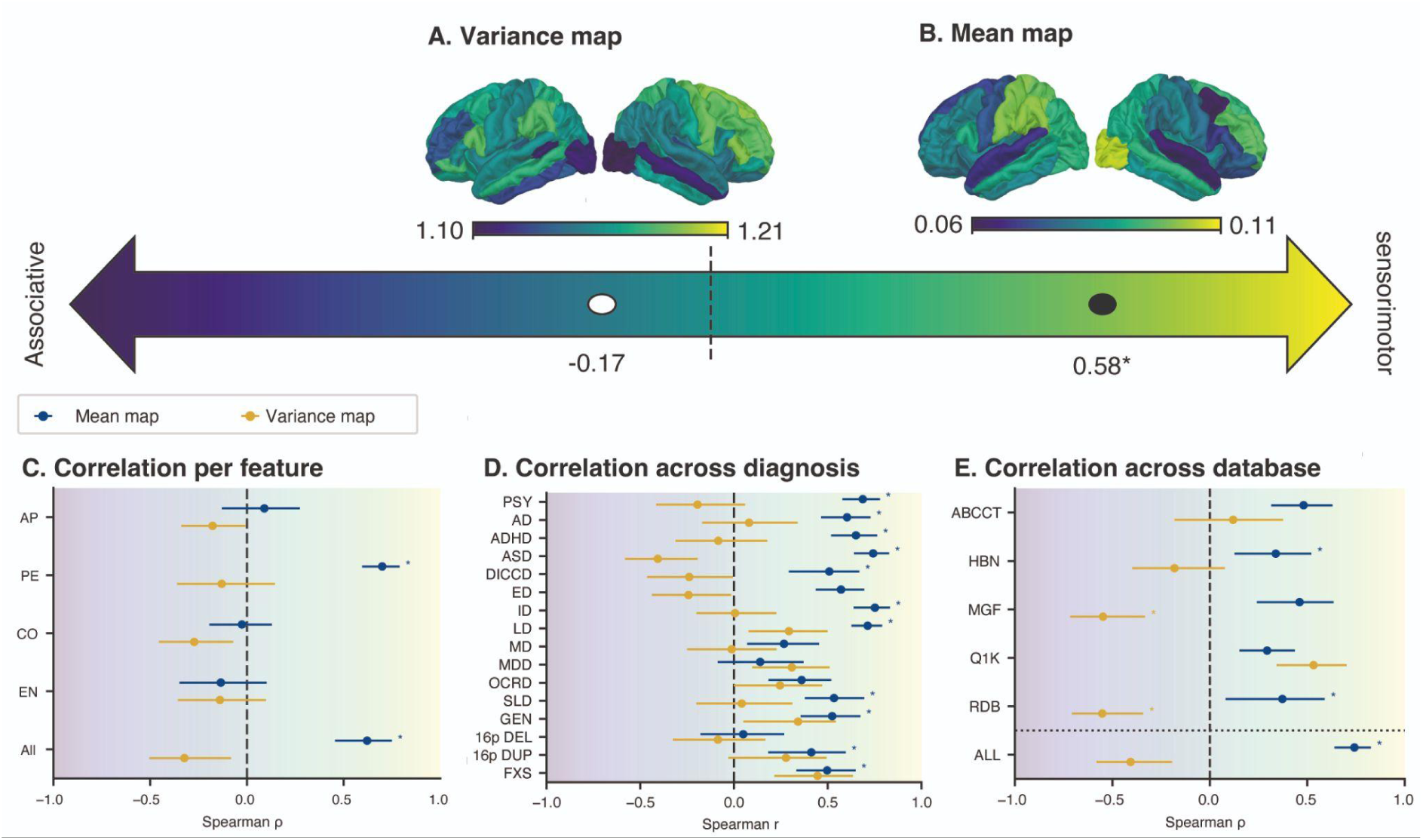
Spatial organization of EEG deviations across cortical hierarchies. **A.** Spatial correlation between the cortical variance deviation map and the cortical gene expression gradient. Warmer colors indicate stronger variance deviations in sensorimotor regions, whereas cooler colors indicate stronger deviations in associative regions. The reported value corresponds to the Spearman correlation coefficient (ρ) between the cortical map and the genetic gradient. **B.** Spatial correlation between the cortical mean deviation map and the genetic gradient. Positive correlations indicate that deviations are preferentially localized in sensorimotor regions. Asterisks indicate significance based on spin permutation testing (q < 0.05). **C.** Spearman correlations between cortical deviation maps and the genetic gradient computed separately for each EEG feature group. Blue points represent mean deviation maps and yellow points represent variance deviation maps. Error bars indicate 95% confidence intervals across diagnoses. **D.** Spearman correlations between cortical deviation maps and the genetic gradient computed separately for each diagnosis. Blue points represent mean deviation maps and yellow points represent variance deviation maps. Error bars indicate 95% confidence intervals across EEG feature groups. **E.** Spearman correlations between cortical deviation maps of Autism Spectrum Disorder (ASD) and the genetic gradient computed separately for each database. Blue points represent mean deviation maps and yellow points represent variance deviation maps. Error bars indicate 95% confidence intervals across diagnoses. The dashed vertical line indicates zero correlation. Background shading illustrates the associative-to-sensorimotor axis of the cortical hierarchy. Abbreviations: ABCCT: Autism Biomarkers Consortium for Clinical Trials, AD: Anxiety Disorder, ADHD: Attention-Deficit/Hyperactivity Disorder, ap: aperiodic, ASD: Autism Spectrum Disorder, co: connectivity, DEL: deletion, DICCD: Disruptive, Impulse-Control, and Conduct Disorders, DUP: duplication, ED: Elimination Disorders, en: entropy, FXS: Fragile X Syndrome, GEN: genetic, HBN: Healthy Brain Network, ID: Intellectual Disability, LD: Language Disorder, MD: Motor Disorders, MDD: Major Depressive Disorder, MGF: Montreal Genetic Family, OCRD: Obsessive-Compulsive and Related Disorders, pe: periodic, PSY: psychiatric conditions, Q1K: Quebec 1000, RDB: Robert Debré University Hospital Center, SLD: Specific Learning Disorder, TD: Typically Developing, UNIGE: Université de Genève, 16p DEL: 16p11.2 deletion, 16p DUP: 16p11.2 duplication.

### Feature wide EEG associations with psychiatric conditions require hundreds of individuals

We tested the sample size required to achieve adequate empirical power using bootstrap resampling (Figure 5A).

**Figure 5.**
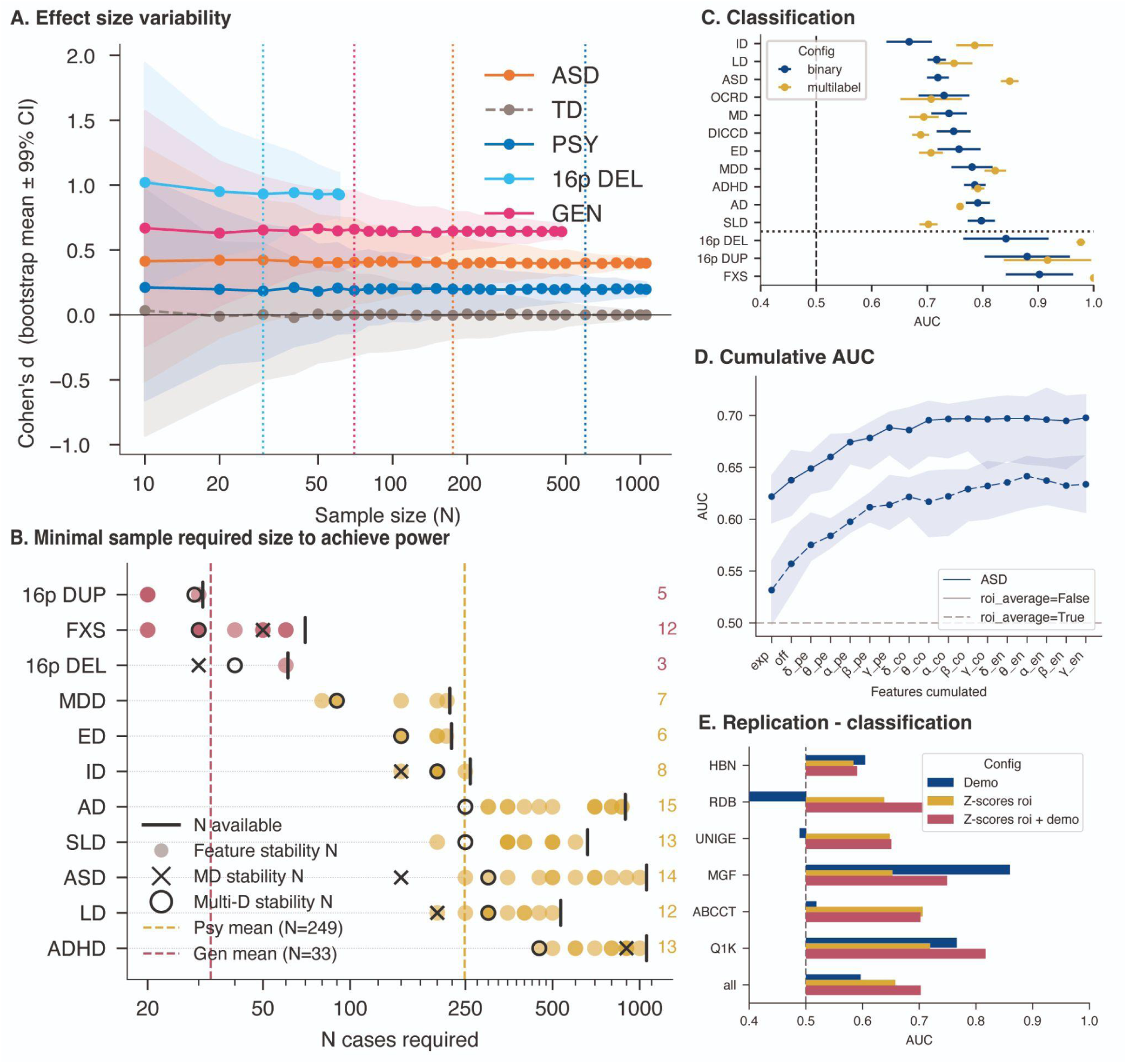
Power, prediction and replication. **A.** Each line shows the bootstrap mean Cohen’s d (± 95% CI, shaded) of the mahalanobis distance as a function of case sample size N (log scale), computed from n=500 bootstrap iterations with balanced cases-typically developing (TD) subsampling. The vertical dotted line marks the minimum N at which empirical power first reaches 95 %. **B.** Each row corresponds to a diagnosis, sorted by the minimum sample size required to achieve stable detection across any feature. The vertical black tick indicates the number of available cases (N available) for that diagnosis, capped at the number of TD. Small dots show the N required for each individual EEG feature to reach empirical power (95%); color indicates diagnosis category (yellow = psychiatric conditions, red = genomic predispositions). The black × marks the N required for the Mahalanobis Distance. The open circle marks the N required for the multivariate effect size across all EEG features simultaneously. The number to the right of each row is the count of features with a valid stability estimate. The x-axis is log-scaled; vertical dashed lines indicate reference sample sizes. **C.** Classification performance (AUC) across diagnoses comparing binary (cases versus TD) and multilabel (all cases simultaneously) prediction models. Points indicate mean AUC values and error bars indicate variability across cross-validation folds. The vertical dashed line indicates chance-level performance (AUC = 0.5). **D.** Classification performance across EEG feature sets for ASD. Solid and dashed lines compare models computed with and without ROI averaging, respectively. Shaded regions indicate variability across cross-validation folds. **E.** Database generalization of ASD classification across feature configurations. Each row represents a held-out database (leave-one-database-out cross-validation). Bar length indicates AUC when training on all remaining databases and testing on the held-out database. The ‘all’ row shows AUC across all databases. Chance level at AUC = 0.5 (dashed line). Databases are ranked by increasing AUC on z-scores. Abbreviations: ABCCT: Autism Biomarkers Consortium for Clinical Trials, AD: Anxiety Disorder, ADHD: Attention-Deficit/Hyperactivity Disorder, ap: aperiodic, ASD: Autism Spectrum Disorder, co: connectivity, DEL: deletion, DICCD: Disruptive, Impulse-Control, and Conduct Disorders, DUP: duplication, ED: Elimination Disorders, en: entropy, FXS: Fragile X Syndrome, GEN: genetic, HBN: Healthy Brain Network, ID: Intellectual Disability, LD: Language Disorder, MD: Motor Disorders, MDD: Major Depressive Disorder, MGF: Montreal Genetic Family, OCRD: Obsessive-Compulsive and Related Disorders, pe: periodic, PSY: psychiatric conditions, Q1K: Quebec 1000, RDB: Robert Debré University Hospital Center, SLD: Specific Learning Disorder, TD: Typically Developing, UNIGE: Université de Genève, 16p DEL: 16p11.2 deletion, 16p DUP: 16p11.2 duplication.

For psychiatric conditions, analyses based on multivariate EEG features required (Figure 5B), on average, approximately 498 participants (249 cases and 249 TD). This varied across diagnoses and sample size was positively and negatively correlated with variance and effect size respectively (i.e., for ASD, we estimated a sample size of n=600). Overall, multivariate Cohen’s d provided improved power compared to individual features. Mahalanobis distance provided the best power to detect deviation in conditions with putative large but heterogeneous effects (e.g., ID, ASD and disruptive disorder). Single genetic variant groups required approximately eight times fewer participants, with 66 participants (33 cases and 33 TD) due to large and homogeneous effect sizes.

### Multifeature approaches predict psychiatric conditions

We assessed the predictive performance of EEG deviations by training a Random Forest classifier on individual-level z-scores. In binary classification, normative EEG deviations discriminated between each diagnostic group and TD with an AUC (cross-validated) ranging from 0.69 to 0.8 for psychiatric conditions and above 0.8 for single genetic variants such as FXS and 16p DUP (Figure 5C). We further evaluated the cumulative contribution of additional features and observed an increase of 0.10 in AUC as features were progressively added, both at the ROI level and at the ROI-averaged level (Figure 5D). We evaluated the generalizability of classification performance by training and testing our model on independent datasets. The mean AUC was 0.72 and 0.66 (Figure 5C) with and without demographic information highlighting the known age and sex biases in psychiatric conditions.

## Discussion

We assembled the largest high-density resting-state EEG datasets to date, spanning 4,812 participants, 11 psychiatric conditions and rare genetic variants. Using normative models, we characterized developmental trajectories of EEG features at the source level in TD participants.

Most developmental changes occurred from 0 to 20 years old, corroborating previous work linking developmental changes of the spectral activity to biological processes^37^. Importantly, the present findings are derived from source space and largely replicate prior scalp-level developmental EEG literature, including age-related increases in alpha center frequency^38^, periodic alpha power^28,39^, alpha-connectivity^40^, and broadband complexity^41^, as well as decreases in aperiodic components^28,37^, suggesting that these developmental trajectories are robust across spatial scales. The notable exception is gamma periodic power, which decreased with age in source space, contrasting with Mallus et al.^28^ who report increasing gamma at the scalp level after similar aperiodic decomposition. Beyond alpha-band connectivity and broadband entropy, no prior normative benchmarks exist; the trajectories reported here therefore provide a reference for future mechanistic and clinical investigations.

Deviations in psychiatric conditions were small and highly correlated. In contrast, individual genetic variants showed effect sizes two- to four-fold larger and strong anti-correlations, extending previous reports of marked neurobiological alterations associated with CNVs^20,42^. The 16p11.2 duplication and deletion variants showed opposing deviation patterns, consistent with previous reports of mirror effects in functional connectivity^43^ and EEG task^44^. When genetic variants were pooled, effect sizes decreased to levels comparable to psychiatric conditions, supporting the presence of opposing variant-specific effects. The shared deviation profile observed across psychiatric conditions and pooled genetic predisposition groups suggests convergence toward common electrophysiological alterations across distinct etiologies. In psychiatric conditions, this convergence is consistent with dimensional models of psychopathology, which posit a shared general factor underlying diagnostically distinct conditions^45,46^, while common environmental and socio-cultural influences associated with living with a condition may also contribute^47^. Groups with greater EEG feature variability, such as ASD and ID, as well as groups pooling different genetic variants, tended to show the smallest effect sizes. For those conditions with more heterogeneity, using the Mahalanobis distance revealed bigger deviations from TD.

Several mechanisms may contribute to these shared electrophysiological alterations. Deviations in periodic power followed patterns consistent with delayed cortical maturation relative to normative developmental trajectories. However, delayed maturation alone cannot account for the full spectrum of observed alterations, such as those observed in aperiodic components or gamma entropy. The aperiodic exponent has been proposed as a marker of excitation–inhibition balance, with a flatter slope reflecting a shift toward relative excitation^48,49^, suggesting that altered E/I dynamics may represent an additional mechanism. At the anatomical level, convergent thalamic alterations across psychiatric conditions have been reported^43^. This convergence is of particular interest given the central role of thalamocortical circuits in generating and coordinating cortical oscillations across frequency bands^50,51^.

The spatial organization of EEG deviations was not uniformly distributed across the cortex but aligned with a cortical gene expression gradient (PC1 gene expression), spanning primary sensorimotor to higher-order association regions. Prior work has shown similar alignment between electrophysiological dynamics and gene expression gradients in typical populations^52–54^. Here, we observed that deviations primarily affect sensorimotor regions. It has been shown that at the anatomical level, genetic variants and psychiatric conditions correlate with this gradient but in opposite directions, with genetic variants preferentially affecting sensorimotor regions and psychiatric conditions more strongly involving association cortices^55^. However, we did observe a convergence on a similar sensorimotor-dominant spatial pattern in the electrophysiological domain.

Historically, EEG studies of psychiatric conditions have often relied on small samples, with approximately 30 cases and TD being common in earlier work^56^. Recent studies with larger datasets highlighted the failure to replicate and the inflation of effect sizes^57,58^ as well as the substantial need for larger cohorts to achieve empirical power^58^. Our analyses corroborate this: multivariate features in psychiatric conditions required large sample sizes to achieve empirical power, with hundreds of participants (cases and controls) on average. In contrast, individual genetic predisposition groups showed larger effect sizes and required substantially fewer participants, with reliable effects achieved with fewer than a hundred participants. Conditions with more variance, such as ASD, benefit from using the Mahalanobis distance rather than group effects. Finally, deviation z-scores showed consistent replication across databases and provided discriminative value for classification analyses.

Several limitations should be acknowledged. While source reconstruction was validated against an independent magnetoencephalography (MEG) atlas, volume conduction effects cannot be fully eliminated, potentially introducing spurious connectivity estimates^59^. Several diagnostic groups and genetic variants had limited sample sizes, reducing statistical power and the reliability of group-level estimates for these conditions. Furthermore, with the exception of ASD, most diagnostic groups are predominantly drawn from a single database (HBN), which limits the assessment of cross-site replicability for these conditions and may introduce site-specific confounds in their deviation profiles. The normative models were trained on TD individuals drawn from a specific age range and demographic background, which may limit generalizability to populations not represented in the training data. Additionally, diagnostic records were inconsistent across datasets, as participants were recruited on the basis of a primary diagnosis or genetic variant without comprehensive screening for co-occurring conditions, potentially underestimating true diagnostic co-occurrence and limiting the interpretation of group-specific deviation profiles. Diagnostic ascertainment also varied in rigor across cohorts: gold-standard research instruments (ADOS and ADI-R) were administered in some datasets (ABCCT, CHUV, RDB, UNIGE) or based on a standardized clinical interview with gold-standard confirmation available only for a subset of cases (HBN, KSADS-COMP). This heterogeneity in diagnostic procedures on top of recruitment context (hospital vs research program) may contribute to variability in the ASD deviation profiles across sites.

Across the largest transdiagnostic source-space EEG dataset assembled to date, we show that the apparent weakness of psychiatric neurophysiological signatures emerges from biological aggregation: genetically defined populations exhibit large and sometimes opposing deviations that progressively cancel when heterogeneous etiologies are combined, ultimately reproducing the small, shared effects observed across behavioral diagnoses. These results lay the groundwork for precision medicine approaches to psychiatry in which individual-level electrophysiological profiles, interpreted against lifespan norms, could inform diagnosis, subgroup stratification, and the evaluation of targeted interventions. A key next step will be to translate these normative profiles from research-grade recordings to the clinical EEG routinely acquired in hospital settings, where lower channel counts, shorter recordings, and heterogeneous acquisition conditions pose challenges that must be addressed before deviation-based metrics can be deployed in practice.

## Methods

### Data

The combined sample included typically developing individuals (TD; *n* = 1,082), individuals with psychiatric conditions (PSY; *n* = 3,329), and individuals with genetic predisposition variants (GEN; *n* = 483). Participants in the GEN group were further classified as carrying deletions (DEL; *n* = 241), duplications (DUP; *n* = 142), or single-nucleotide variants (SNVs; *n* = 34). For specific genetic variants with sufficient sample sizes (*n* ≥ 30), we also performed separate analyses. These included fragile X syndrome (FXS), 16p11.2 deletion (16p DEL; *n* = 61), and 16p11.2 duplication (16p DUP; *n* = 31). Note that the PSY, DEL, DUP, SNV, and GEN groups are not mutually exclusive; some participants in the GEN group also carried psychiatric diagnoses. However, most participants with a genetic predisposition showed minimal co-occurrence with psychiatric diagnoses, consistent with their smaller sample sizes and the ascertainment bias of each dataset, in which participants were recruited either on the basis of their genetic variant or their psychiatric diagnosis, but rarely both. Sample sizes, demographic characteristics, and dataset provenance are summarized in Table 1. See Table S1 for the complete list of genetic variants.

**Table 1.** EEG cohorts stratified by diagnoses and genetic variants.

| label | N | sex (M/F) | age | HBN | MGF | RDB | Q1K | ABCCT | UNIGE | CIN | SL | CHUV |
| --- | --- | --- | --- | --- | --- | --- | --- | --- | --- | --- | --- | --- |
| Unselected/general population |  |  |  |  |  |  |  |  |  |  |  |  |
| TD | 1082 | 562/520 | 19±16 | 227 | 279 | 77 | 160 | 123 | 112 | 71 | 14 | 19 |
| Ascertained for psychiatric conditions |  |  |  |  |  |  |  |  |  |  |  |  |
| ADHD | 1766 | 1209/557 | 11±5 | 1468 | 64 | 117 | 117 | 0 | 0 | 0 | 0 | 0 |
| ASD | 1309 | 988/321 | 10±5 | 351 | 55 | 335 | 116 | 219 | 233 | 0 | 0 | 0 |
| AD | 904 | 501/403 | 12±6 | 826 | 25 | 0 | 53 | 0 | 0 | 0 | 0 | 0 |
| SLD | 668 | 416/252 | 11±5 | 603 | 23 | 0 | 42 | 0 | 0 | 0 | 0 | 0 |
| LD | 541 | 361/180 | 10±5 | 320 | 122 | 0 | 99 | 0 | 0 | 0 | 0 | 0 |
| DICCD | 507 | 319/188 | 10±3 | 384 | 3 | 92 | 28 | 0 | 0 | 0 | 0 | 0 |
| ID | 264 | 166/98 | 14±11 | 91 | 107 | 10 | 42 | 14 | 0 | 0 | 0 | 0 |
| MDD | 227 | 111/116 | 14±5 | 221 | 6 | 0 | 0 | 0 | 0 | 0 | 0 | 0 |
| ED | 227 | 168/59 | 9±3 | 227 | 0 | 0 | 0 | 0 | 0 | 0 | 0 | 0 |
| MD | 223 | 177/46 | 10±4 | 200 | 23 | 0 | 0 | 0 | 0 | 0 | 0 | 0 |
| OCRD | 121 | 65/56 | 12±4 | 118 | 3 | 0 | 0 | 0 | 0 | 0 | 0 | 0 |
| PSY | 3329 | 2232/1097 | 11±6 | 2111 | 208 | 357 | 201 | 219 | 233 | 0 | 0 | 0 |
| Ascertained for genetic liability |  |  |  |  |  |  |  |  |  |  |  |  |
| FXS | 70 | 38/32 | 21±10 | 0 | 0 | 0 | 0 | 0 | 0 | 70 | 0 | 0 |
| 16p DEL | 61 | 31/30 | 11±13 | 0 | 23 | 0 | 0 | 0 | 0 | 0 | 25 | 13 |
| SNV | 34 | 21/13 | 11±9 | 0 | 12 | 0 | 22 | 0 | 0 | 0 | 0 | 0 |
| 16p DUP | 31 | 23/8 | 12±14 | 0 | 15 | 0 | 1 | 0 | 0 | 0 | 15 | 0 |
| DUP | 142 | 89/53 | 16±15 | 0 | 103 | 0 | 22 | 0 | 0 | 0 | 15 | 2 |
| DEL | 241 | 120/121 | 14±14 | 0 | 156 | 0 | 44 | 0 | 0 | 0 | 25 | 16 |
| GEN | 483 | 265/218 | 16±14 | 0 | 270 | 0 | 85 | 0 | 0 | 70 | 40 | 18 |
| TOTAL | 4812 | 3014/1798 | 13±11 | 2404 | 598 | 515 | 376 | 345 | 342 | 141 | 54 | 37 |
| ABCCT: Autism Biomarkers Consortium for Clinical Trials, CIN: Cincinnati Children's Hospital, CHUV: Centre Hospitalier Universitaire Vaudois, HBN: Healthy Brain Network, Q1K: Quebec 1000, MGF: Montreal Genetic Family, UNIGE: Université de Genève, RDB: Robert Debré University Hospital Center, SL: Simons Searchlight. AD: Anxiety Disorder, ADHD: Attention-Deficit/Hyperactivity Disorder, ASD: Autism Spectrum Disorder, DICCD: Disruptive, Impulse-Control, and Conduct Disorders, ED: Elimination Disorders, ID: Intellectual Disability, LD: Language Disorder, MDD: Major Depressive Disorder, MD: Motor Disorders, OCRD: Obsessive-Compulsive and Related Disorders, PSY: psychiatric conditions, SLD: Specific Learning Disorder, TD: Typically Developing. |  |  |  |  |  |  |  |  |  |  |  |  |

**DEL:** deletion, **DUP:** duplication, **FXS:** fragile X syndrome, **GEN:** genetic, **16p DEL:** 16p11.2 deletion, **16p DUP:** 16p11.2 duplication

All datasets were acquired using 128-channel HydroCel Geodesic Sensor Nets (Magstim EGI, Eugene, OR, USA) under eyes-open resting-state conditions. Sampling rates ranged from 250 to 1000 Hz across sites, with Cz as the online reference and impedances maintained below site-specific thresholds (10–50 kΩ). Diagnoses were established by licensed clinicians according to DSM-5 criteria. Genetic diagnoses were confirmed through chromosomal microarray analysis (CMA) for CNV carriers, or Southern blot and PCR testing for FXS participants (CGG triplet repeat expansions exceeding 200 repeats in the FMR1 gene). Site-specific acquisition parameters, diagnostic procedures, and data access information are provided in Supplementary Information “Data acquisition”.

### EEG signal processing

#### Preprocessing

We developed a Python script using the MNE package60 for automatic preprocessing of raw data following these steps: (1) Apply a notch filter at 60 Hz and its harmonics (North American datasets) or 50 Hz and its harmonics (European datasets) to remove line noise; (2) Resample the raw data to 250 Hz; (3) Remove flat channels with a -3 SD threshold applied to the log-transformed standard deviation of each channel; (4) Set average reference and apply the GSN-HydroCel-128 montage; (5) Apply ICA (Independent Component Analysis) using the Infomax algorithm with 25 components to identify and remove artifactual components, including eye movements, muscle activity, and other non-brain sources, using the ICLabel classifier^61^; (6) Apply a bandpass filter between 0.1 and 48 Hz; (7) Segment continuous EEG into 2-s non-overlapping epochs; (8) Remove bad segments and interpolate artifactual channels within each epoch using the Autoreject library with Bayesian optimization and up to 32 interpolated channels^62^.

#### Source reconstruction

Cortical sources were reconstructed from preprocessed EEG epochs using the eLORETA method^63^ implemented in MNE-Python^60^. Age-appropriate head models were used: for participants older than 3 years, the fsaverage template from FreeSurfer^64^ was used with an identity transformation, while younger participants were processed using age-specific anatomical templates from the ANTS pediatric atlas, with digitized electrode montages aligned in native space. A predefined set of reference electrodes known to carry limited neural signals was excluded prior to source reconstruction. A forward solution was computed using a three-layer boundary element model with a minimum source distance of 5 mm, and noise covariance was estimated from the preprocessed epochs. An inverse operator was constructed using a regularization parameter of lambda2 = 1/9 (corresponding to an assumed signal-to-noise ratio of 3), and eLORETA was applied to each epoch with normal orientation. Regional time series were extracted by averaging source activity within cortical regions defined by the Desikan-Killiany atlas^65^, yielding 68 regional time series for subsequent analyses.

### EEG features

We extracted spectral, connectivity, and entropy features from cortical ROIs defined by the parcellation described above (see EEG preprocessing). Spectral features were decomposed into aperiodic components (offset and exponent) and periodic components, comprising power across canonical frequency bands (delta: 2-4 Hz, theta: 4-8 Hz, alpha: 8-12 Hz, beta: 12-30 Hz, gamma: 30-45 Hz) and peak characteristics (center frequency, peak power, and bandwidth). Connectivity and entropy were extracted across the same canonical frequency bands. In total, p = 20 EEG features (comprising 2 aperiodic, 5 periodic, 3 peak, 5 connectivity, and 5 entropy features) were used for multivariate statistical analyses. When using the region of interest, we used 17 features × 68 ROIs = 1156 features. The 3 features characterizing the peak are not used at the region level, as they are not properly characterized in regions far from the occipital.

#### Power spectral density

PSD was computed using Welch’s method66, applied to 2-second sliding windows with 50% overlap and Hamming tapering. PSD values were log-transformed for normalization and averaged across epochs.

#### Periodic power and aperiodic components

The periodic power and aperiodic components were parameterized using the SpecParam algorithm^48,67^ with the following settings: peak width limits = [1, 8], maximum number of peaks = 6, minimum peak height = 0.1, peak threshold = 2, and aperiodic mode = ‘fixed’. The aperiodic component was first estimated across the full spectrum and subtracted to isolate periodic components. Aperiodic parameters, including the exponent and offset, were extracted for further analysis. Periodic spectral fits revealed increased delta power in frontal regions, partly reflecting a low-frequency bend in the source-reconstructed power spectra (see Supplementary Information, *EEG decomposition*)

#### Functional connectivity analysis

Source-level functional connectivity was estimated using the weighted phase lag index^68^ using the time-resolved spectral connectivity implementation in MNE- Connectivity (mne_connectivity.spectral_connectivity_time;^69^). For each ROI, we computed the mean wPLI across all connections to other ROIs, providing a summary measure of each region’s overall connectivity strength within each frequency band. This approach identifies hub regions with elevated network connectivity.

#### Entropy

Band-specific permutation entropy was computed to quantify the temporal complexity of neural oscillations within each frequency band^70^, using the *AntroPy* Python library^71^. PE was calculated for each epoch and ROI using an embedding dimension of 3 and a time delay of 1 sample, and normalized by dividing by log₂(3!) to yield values between 0 and 1, where higher values indicate greater signal complexity. Values were then averaged across epochs for each frequency band.

#### Mahalanobis distance

Mahalanobis distance was used to quantify the multivariate distance of each subject from the typically developing (TD) control group. The control-group mean vector and covariance matrix were estimated from the control subjects, defining the reference centroid and covariance structure. For each subject, the Mahalanobis distance was calculated as: D(x) = √[(x − μ)ᵀ S⁻¹ (x − μ)] where x is the subject’s feature vector, μ is the control-group mean vector, and S is the control-group covariance matrix. Mean imputation was applied only at the scoring stage, when necessary, to obtain a distance for subjects with incomplete feature vectors and was not used to estimate the control-group mean or covariance matrix.

#### Multivariate Cohen’s d

Multivariate Cohen’s d was used to quantify the magnitude of multivariate separation between each diagnostic group and the TD. The squared effect size was calculated as the squared Mahalanobis distance between the two group centroids: D² = δᵀ Σw⁻¹ δ, where δ = x̄g − x̄c represents the difference between the diagnostic-group and control-group mean vectors, and Σw is the pooled within-group covariance matrix. The multivariate effect size was then calculated as D = √D². Group means and the pooled covariance matrix were estimated from the same complete-case observations; therefore, ng and nc refer to the complete-case sample sizes of the diagnostic and control groups, respectively. Because D² is positively biased in finite samples, a bias correction was applied. With N = ng + nc, the corrected squared distance was calculated as: D²corrected = [(N − p − 3) / (N − 2)] × D² − p × (1/ng + 1/nc). The corrected effect size was then calculated as Dcorrected = √max(D²corrected, 0). This correction accounts for both the multiplicative and additive components of the finite-sample bias.

#### Generalized variance ratio

This measure was used to quantify differences in overall multivariate dispersion between each diagnostic group and the TD. For each group, generalized variance was defined using the pseudo-determinant of its covariance matrix, calculated as the product of its strictly positive eigenvalues (threshold = 1 × 10⁻¹⁰). This approach accommodates rank-deficient covariance matrices when the number of features approaches or exceeds the available group sample size. The GVR was calculated as the difference between the bias-corrected log pseudo-determinants of the diagnostic-group and control-group covariance matrices: V = log det⁺(Sg) − log det⁺(Sc), where Sg and Sc are the covariance matrices of the diagnostic and control groups, respectively. Positive V values indicate greater overall multivariate dispersion in the diagnostic group, whereas negative values indicate lower dispersion relative to TD. To account for the small-sample bias of the log-determinant, a bias correction based on the expected log-determinant of the sample covariance matrix under the Wishart distribution was applied before calculating V. The expected value was defined as E[log det(S)] = log det(Σ) + Σᵢ[ψ((n − i)/2) − log((n − 1)/2)], where ψ is the digamma function, n is the group sample size, and i ranges from 1 to min(p, n − 1). The correction accounts for the dependence of the log-determinant bias on sample size and dimensionality, thereby reducing systematic differences between groups arising from unequal sample sizes.

### Normative modeling

Given the wide age range of participants and the non-linear nature of brain development across childhood and adolescence, spectral and connectivity features were adjusted using a normative modeling approach implemented with the PyNM package (v1.0.0b8; https://github.com/ppsp-team/PyNM)^72^. Normative modeling accounts for developmental variability by modeling individual deviations relative to a neurotypical reference population^73,74^. In this study, the GAMLSS was used to model typical developmental trajectories using a four-parameter distribution such as Sinh-Arcsinh ^75^, where the location (µ, representing the mean) and scale (σ, representing the variance) are modeled as functions of age, sex, and site, and shape parameters (ν for skewness and τ for kurtosis) were included to account for the distributional properties of the data.

EEG features were used as the dependent variable in the normative model. Outliers were identified using an interquartile range (IQR) method computed on all observations: observations falling below the 15th percentile minus 1.5×IQR or above the 85th percentile plus 1.5×IQR were excluded to mitigate the influence of extreme values. Covariates included age, sex, recording site, and data quality, with site treated categorically as a fixed effect to account for potential batch effects, and data quality defined as the ratio of bad segments identified via Autoreject^62^. The normative model was fitted at two levels of spatial resolution depending on the analysis: features were averaged across ROIs when quantifying multivariate deviations across diagnostic groups; and models were fitted separately for each ROI when mapping the spatial organization of deviations along cortical gene expression gradients.

#### Trajectories

When visualizing developmental trajectories, the recording site was included as a fixed effect in the GAMLSS model, with the first site alphabetically serving as the reference category. To center trajectories on a site-neutral reference, the estimated site offset of the mode database was subtracted from individual predictions prior to plotting. Trajectories were computed separately for males and females and displayed as smooth fitted curves with 95% confidence intervals, overlaid on the site-corrected scatter of individual data points from typically developing and clinical participants. All lifespan trajectories at the ROI level, along with the corresponding model quality metrics, are available at https://dub21.github.io/D-EEG/

#### Model performance

For each model, we assessed the normality, skewness, and kurtosis of the group’s residuals. Additionally, we computed the Akaike Information Criterion (AIC) and the Bayesian Information Criterion (BIC)^76^ to compare model fit. A detailed table presenting these statistics for each ROI across all frequency bands is provided at https://dub21.github.io/D-EEG.

#### Statistical analysis

To compare diagnostic groups against the TD group, the mean z-score derived from the normative model was tested for each group against the TD group using a permutation test. Multiple comparisons were corrected using the Benjamini-Hochberg false discovery rate (FDR) procedure, with statistical significance set at FDR-corrected q < 0.05. Effect sizes were expressed as the Cohen’s d.

### Replicability and stability analysis

#### Empirical power

To estimate empirical power as a function of sample size, we performed a subsampling analysis for each feature and diagnosis. Subsamples were drawn without replacement at linearly spaced sample sizes: from 10 to 100 in steps of 10, from 125 to 250 in steps of 25, from 300 to 500 in steps of 50, from 500 to the maximum available sample size in steps of 100, plus the maximum available sample size itself, with 500 iterations per step. At each iteration, an equal number of TD was drawn (without replacement) to match the case subsample size. For univariate features, statistical significance was assessed using Welch’s t-test. For the Mahalanobis distance, significance was assessed using the Mann-Whitney U test. For the multivariate effect size, significance was assessed using Hotelling’s T², converted to an F-test. These tests were applied consistently across all sample sizes. Empirical power at each sample size was defined as the proportion of iterations yielding a significant result (p < 0.05). The minimum sample size required to reach 95% empirical power was reported, provided the effect was also significant at the full sample size. At this power level, the Type II error rate (β = 1 − power = 5%) is equivalent to the Type I error rate (α = 5%), balancing the two types of statistical error.

#### Feature replication

To assess the cross-dataset generalizability of EEG deviations, we computed a leave-one-out (LOO) sign replication rate for each combination of feature and acquisition site. For each focal database, Cohen’s d was computed between the diagnostic group and TD within that database, and independently on all remaining databases pooled (the LOO reference). A replication was counted when the sign of the focal effect matched the sign of the LOO reference. This yielded a database × features binary matrix of sign agreements (0 or 100%).

### Cortical gradients

We used previously published normative maps of cortical organization hierarchies^32^, specifically the spatial transcriptomic map (PC1 gene expression)^36,77^. To assess spatial correlations with this normative reference, all maps were projected onto the 68 cortical regions of the Desikan parcellation using the *neuromaps* Python package^32^. The statistical significance of the spatial correlations was then evaluated using a spin-permutation procedure^78,79^.

### Model prediction

We trained a Random Forest classifier^80^ of 300 decision trees, each built on a bootstrap sample of the training data with square-root feature subsampling at each node. Class imbalance was addressed by setting class weights inversely proportional to class frequencies. A minimum of 5 samples per leaf was enforced to reduce overfitting. Features consisted of normative deviation z-scores averaged across ROIs, yielding one value per EEG feature type and frequency band per participant. Classification performance was evaluated using the area under the receiver operating characteristic curve (AUC-ROC), which is threshold-independent and robust to class imbalance. Model generalization was assessed via stratified 5-fold cross-validation or, alternatively, leave-one-database-out cross-validation, where the model was trained on all datasets except one and evaluated on the held-out dataset, iteratively over all available datasets. This latter scheme provides a stringent test of cross-site generalization. Feature importance was derived from the mean decrease in impurity across trees, aggregated by EEG feature group to identify which signal dimensions most contributed to classification.

## Code Availability

All code used in this study is available at https://dub21.github.io/D-EEG.

## Data Availability

HBN data are publicly available at https://fcon_1000.projects.nitrc.org/indi/cmi_healthy_brain_network/. Simons Searchlight data are available to approved researchers via SFARI Base (https://base.sfari.org/dataset/DS0000050). All other datasets are available upon reasonable request to the corresponding author. Normative trajectories, brain deviation maps, model diagnostics, and z-scores are available at https://dub21.github.io/D-EEG.

## Acknowledgment

This study was undertaken thanks to funding from the Institute for Data Valorization, Montreal, and the Canada First Research Excellence Fund (IVADO; CF00137433 & PRF3) and was enabled in part by support provided by Calcul Québec (www.calculquebec.ca) and the Digital Research Alliance of Canada (www.alliancecan.ca). G.D. was supported by the Fonds de recherche du Québec - Santé (FRQ-S; 2024-2025 - CB - 350516), Natural Sciences and Engineering Research Council of Canada (NSERC; DGECR-2023-00089), and the Brain Canada Foundation (2022 Future Leaders in Canadian Brain Research program). JCM was supported by NIH U19 MH108206 and NIMH R01 MH138485.

We are grateful to all of the families at the participating Simons Searchlight sites as well as the Simons Searchlight Consortium, formerly the Simons VIP Consortium. We appreciate obtaining access to EEG data on SFARI Base. Approved researchers can obtain the Simons Searchlight population dataset described in this study (https://base.sfari.org/dataset/DS0000050) by applying at https://base.sfari.org.

BRH and NC were supported by the Fondation Hoffmann.

Acquisition of EEG data from the Geneva Autism Cohort (UNIGE) was supported by several grants over the years, including by the Swiss National Foundation Synapsy (Grant No. 51NF40–185897), the Swiss National Foundation for Scientific Research (Grant Nos. #163859, #190084, #202235, #212653 to M.S.), the Fondation Privée des Hôpitaux Universitaires de Genève (https://www.fondationhug.org) and by the Fondation Pôle Autisme (https://www.pole-autisme.ch). The funders were not involved in this study and had no role other than to provide financial support.

The data and/or materials and/or registry resources used for this research were made available by the Quebec 1000 families (Q1K) project and through funds from the Fondation Marcelle et Jean Coutu and the Fonds de Recherche du Québec – secteur santé. The authors also wish to thank all families who contributed to the Q1K project.

## Author information

These authors jointly supervised this work: Sébastien Jacquemont, Guillaume Dumas.

## Contributions

GD and SJ conceptualized and developed the project, supervised the analysis, and substantially revised the manuscript. AEED conducted the analysis and prepared the manuscript, figures, and website. All authors read and approved the final manuscript.

## Corresponding author

Adrien EE Dubois

