## Supplementary Information for "Heterogeneity of brain dynamics in genetic and psychiatric conditions"

#### Common nomenclature

**Cohorts**
ABC-CT: Autism Biomarkers Consortium for Clinical Trials · CHUV: Centre Hospitalier Universitaire Vaudois · CIN: Cincinnati Children's Hospital · HBN: Healthy Brain Network · MGF: Montréal Genetic Family · Q1K: Quebec 1000 · RDB: Robert Debré University Hospital Center · SL: Simons Searchlight · UNIGE: Université de Genève

**Psychiatric conditions**
AD: Anxiety Disorder · ADHD: Attention-Deficit/Hyperactivity Disorder · ASD: Autism Spectrum Disorder · DICCD: Disruptive, Impulse-Control, and Conduct Disorders · ED: Elimination Disorders · ID: Intellectual Disability · LD: Language Disorder · MD: Motor Disorders · MDD: Major Depressive Disorder · OCRD: Obsessive-Compulsive and Related Disorders · PSY: Psychiatric Conditions · SLD: Specific Learning Disorder · TD: Typically Developing

**Genetic conditions**
DEL: Deletion · DUP: Duplication · FXS: Fragile X Syndrome · GEN: Genetic (combined group) · SNV: Single-Nucleotide Variant(s) · 16p DEL: 16p11.2 deletion · 16p DUP: 16p11.2 duplication

**EEG features**
ap: aperiodic · pe: periodic · co: connectivity · en: entropy · α: alpha · β: beta · δ: delta · γ: gamma · θ: theta · Bw: bandwidth · Cf: center frequency · Ex: exponent · Of: offset · Pw: peak power · PSD: power spectral density · wPLI: weighted phase lag index

**Methods**
AIC / BIC: Akaike / Bayesian Information Criterion · AUC(-ROC): Area Under the Curve (ROC) · MVD: Multivariate Cohen's d · d: Cohen's d · GVR / V: Generalized Variance Ratio · IQR: interquartile range · LOO: leave-one-out

### Figures

**A)**

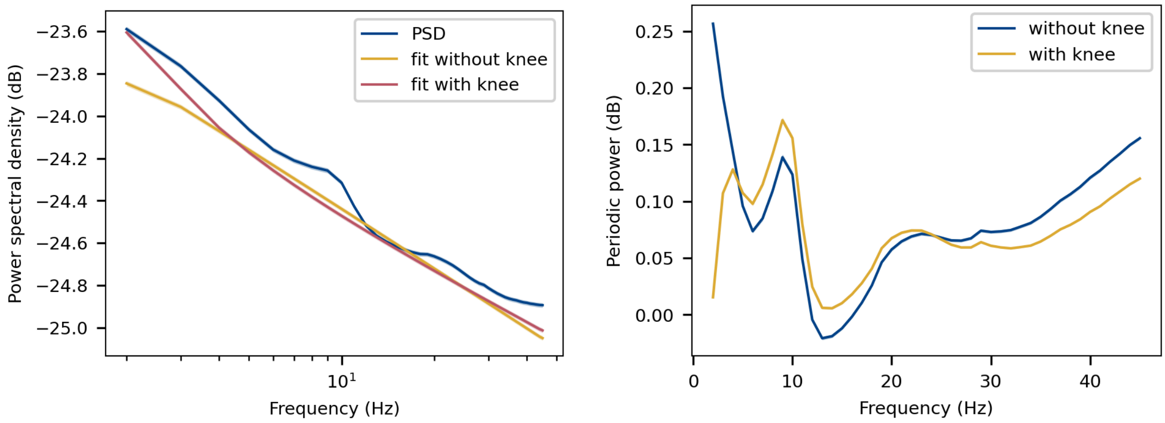


**B)**

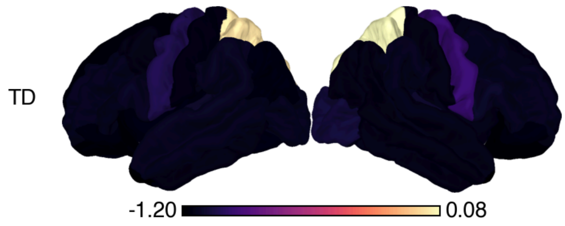


**C)**

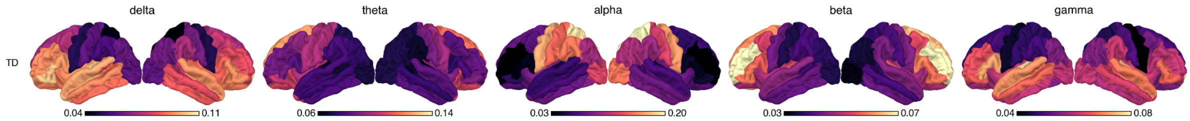


**D)**

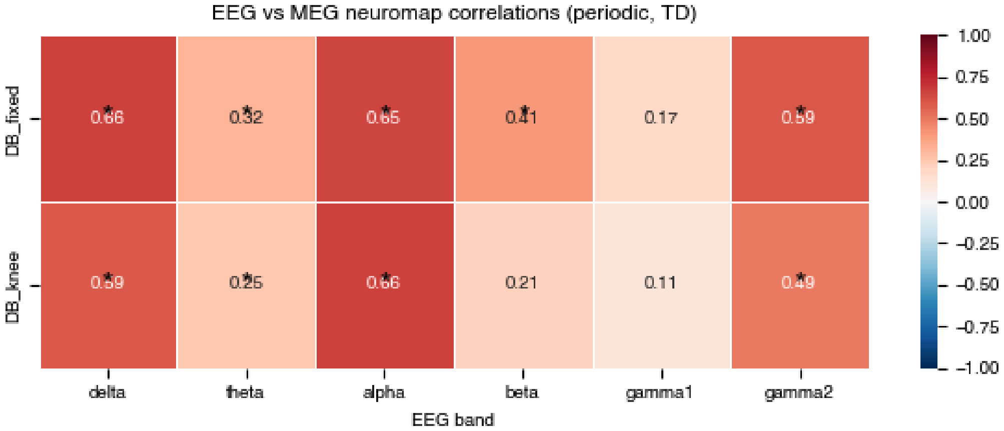

***Figure S1. A)*** *Frequency-dependent spectral characteristics in the typically developing (TD) group. (left) Mean power spectral density (PSD, dB) across frequencies, estimated with and without inclusion of the aperiodic knee in the spectral parameterization. The frequency axis is shown on a logarithmic scale. (right) Mean periodic power (dB) and corresponding aperiodic and periodic model components, estimated with and without inclusion of the aperiodic knee. Lines represent the group mean across participants, and shaded areas indicate the standard error of the mean (SEM). The elevated periodic delta power observed in our analyses could potentially reflect a spectral bend rather than a distinct periodic component. Overall, the sensitivity analyses yielded highly similar results between models with and without a knee. However, the correlation with the cortical gradient indicated that part of the spatial association observed in the original analysis was attributable to the aperiodic knee component.* ***B).*** *Spatial distribution of the aperiodic knee parameter across the cortex. Most estimated knee frequencies fell outside the examined frequency range, limiting the interpretability of this parameter.* ***C)*** *Source-level periodic power (with knee mode) in the typically developing (TD) group across frequency bands: delta (2-4 Hz), theta (4-8 Hz), alpha (8-12 Hz), beta (12-30 Hz) and gamma (30-45 Hz) using the knee parameter.* ***D)*** *Regional correspondence between EEG periodic (oscillatory) power and MEG-derived functional neuromaps across frequency bands in typically developing individuals. For each Desikan-Killiany cortical region, EEG periodic power was averaged across typically developing participants after removing outlier subjects (values outside median ± 3×IQR per region). Regional EEG values were correlated (Pearson's r) with region-averaged MEG neuromaps (obtained via the neuromaps toolbox) separately for six canonical frequency bands (delta, theta, alpha, beta, low gamma [gamma1], high gamma [gamma2]) and for each of several independent MEG maps; correlations were then averaged across maps within each band. Rows correspond to two approaches for fitting the aperiodic component when extracting periodic power: a fixed model (DB_fixed) and a model including a spectral knee parameter (DB_knee). Cell color and annotated values indicate the mean Pearson r (RdBu_r colormap, range −1 to 1); * represent significant correlation ($p < 0.05$).*

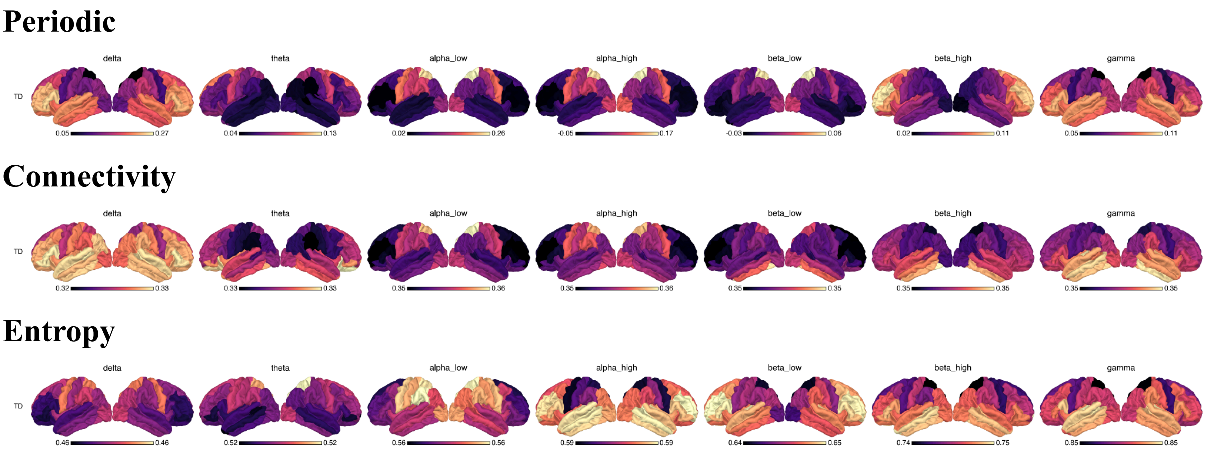


***Figure S2.*** *Source-level periodic power, connectivity, and entropy measures in the typically developing (TD) group across frequency bands. Measures are shown for delta (2-4 Hz), theta (4-8 Hz), low alpha (8-10 Hz), high alpha (10-12 Hz), low beta (12-20 Hz), high beta (20-30 Hz), and gamma (30-45 Hz). Alpha and beta bands were further subdivided into low- and high-frequency components. These sub-bands exhibited distinct topographic transitions: periodic power showed a sharp shift from frontal to occipital dominance across the beta sub-bands, whereas entropy showed a pronounced topographic transition across the alpha sub-bands.*

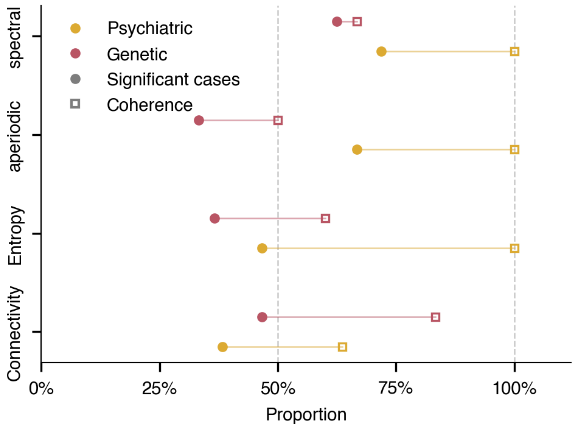


***Figure S3.*** *Each row represents an EEG feature category. For each category, two metrics are shown separately for psychiatric (PSY) and genetic (GEN) groups. Filled circles (●) indicate the proportion of significant feature-diagnosis tests (FDR-corrected < 0.05), assessed via permutation test against the typically developing (TD) reference group. Open squares (□) indicate sign coherence: the proportion of FDR-significant diagnoses showing effects in the same direction relative to TD (Cohen's d sign agreement). The thin line connecting the two markers visualises the gap between significance rate and directional consistency. Feature categories are sorted by proportion of significant tests in descending order. Vertical dashed lines mark 50% (chance level for sign coherence) and 100% (perfect agreement).*

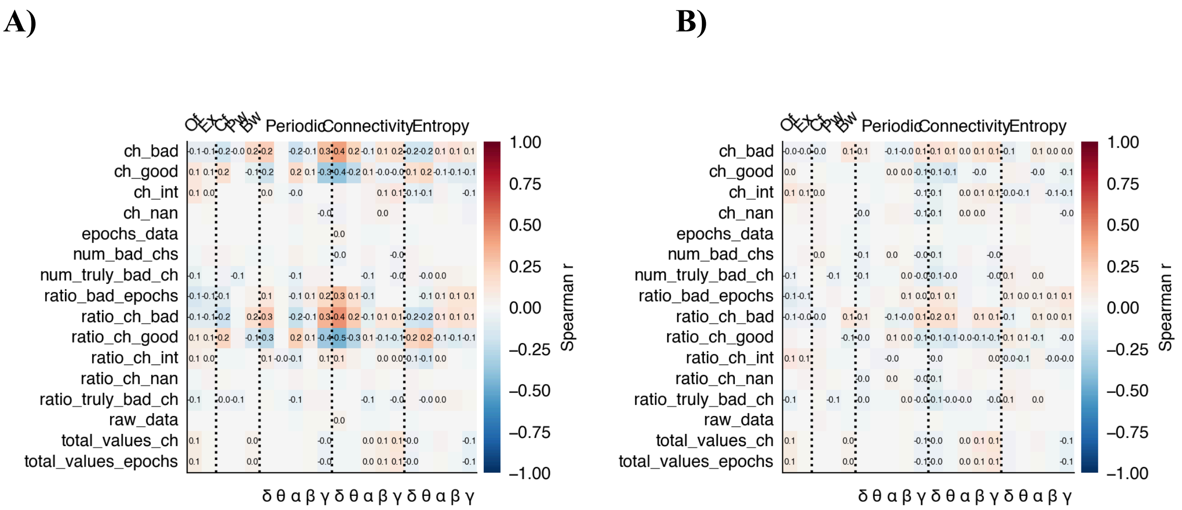


***Figure S4.*** *Each cell shows the Spearman r between the mean z-score (A) without data-quality correction and (B) after data-quality correction across features within the group and the corresponding variable. The numeric value is displayed only when FDR-significant (q < 0.05).*

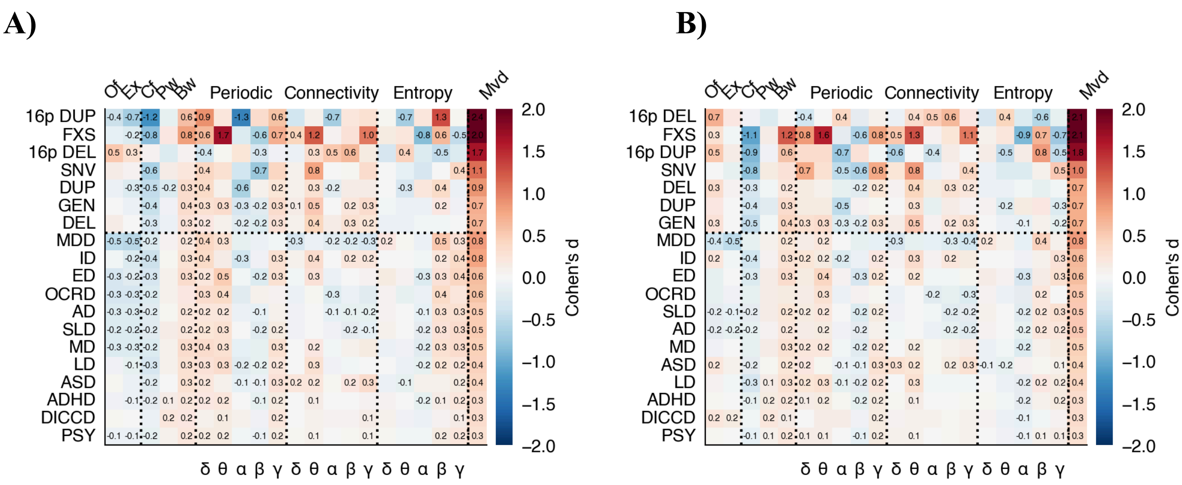


***Figure S5.*** *Sensitivity analyses evaluating the robustness of the main findings to key methodological choices. Each matrix represents a repetition of the main analysis with one parameter modified. (A) Spectral features were extracted using the aperiodic knee parameterization; entropy and connectivity measures were unaffected by this modification. (B) The $R^2$ of the SpecParam model fit was included as a covariate; this analysis was restricted to spectral features. Periodic power, connectivity, and entropy showed consistent patterns across these sensitivity analyses, whereas the aperiodic components (offset and exponent) were more sensitive to model specification.*

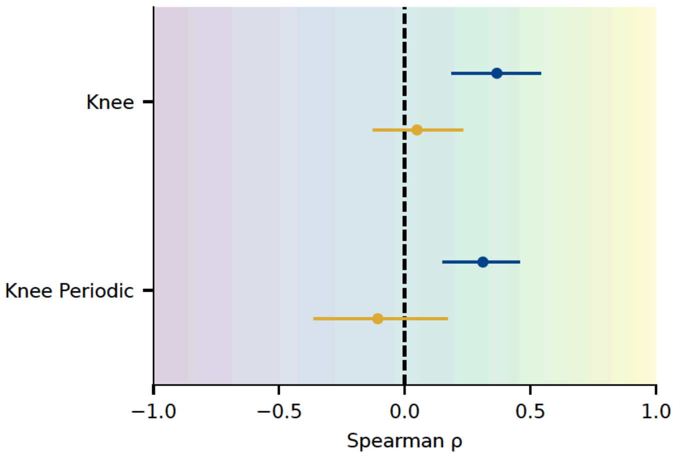


***Figure S6.*** *Spatial correlations between cortical deviation maps and the genetic gradient, computed separately for the aperiodic knee parameter and periodic power estimated using the knee parameterization. Positive correlations indicate that deviations are preferentially localized in sensorimotor regions. Blue points represent correlations with mean deviation maps, whereas yellow points represent correlations with variance deviation maps. Error bars indicate 95% confidence intervals across diagnoses.*

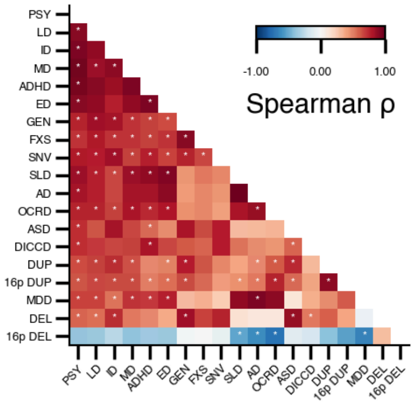


***Figure S7.*** *Each cell represents the Spearman correlation between the mean z-scored EEG feature vectors of two diagnostic groups. Feature vectors were computed as the group mean across all EEG features (averaged across region of interest). Only the lower triangle is shown. Diagnoses are ordered by descending mean correlation across all pairs. *: significant correlation after false discovery rate correction (FDR-BH, α = 0.05; two-sided permutation test). Colour scale ranges from −1 (negative correlation) to +1 (positive correlation).*

### Data acquisition

**ABC-CT**

EEG data were collected across five sites on an EGI 128-channel acquisition system, from the Net Amps 300 series (three sites) or 400 series (two sites), 128-electrode EGI HydroCel Geodesic Sensor Nets, Logitech Z320 computer speakers, a Cedrus StimTracker (for visual presentation timing), and a 23-inch monitor. A standard acquisition setup was implemented: a 1,000-Hz sampling rate, a 0.1- to 200-Hz filter, EGI MFF file format, onset recording of amplifier and impedance calibrations, and a post-acquisition 0.1-Hz digital high-pass filter. E-Prime, version 2.0, was used for experimental control. Cz served as the reference point for all electrodes for recording, and impedances were kept below 50 kΩ. ASD diagnosis was confirmed by the Autism Diagnostic Interview - Revised, Autism Diagnostic Observation Schedule - Second Edition, and clinical confirmation of DSM criteria.

**CHUV**

EEG data were collected using the high-density 128-channel HydroCel geodesic sensor net together with Electrical Geodesics, Inc (Magstim EGI, Eugene, Oregon) Net Station software (v4.5.1). EEG signals were sampled at 250 Hz and bandpass filtered off-line in the range 0.1-100 Hz. Cz electrode was the online recording reference. Impedances were kept below 50 kΩ. All children in the CHUV cohort were referred to the Service des Troubles du Spectre de l'Autisme at Lausanne University Hospital. Trained licensed psychologists and child psychiatrists established formal ASD diagnosis based on the criteria of the Diagnostic and Statistical Manual of Mental Disorders, 5th edition, Text Revision (DSM-5-TR). The clinical procedure for the ASD diagnosis included a review of patients' medical and developmental history as well as the assessment with the Autism Diagnostic Interview - Revised (ADI-R), and the Autism Diagnostic Observation Schedule-2 (ADOS-2).

**CIN**

EEG data were collected using 128-channel EGI HydroCel Geodesic Sensor Nets with a sampling rate of 1000 Hz, referenced to Cz. Five minutes of resting-state EEG were recorded while participants were seated watching a silent video in a quiet environment with eyes open (Liu et al., 2023). Prior to data collection, trained staff familiarized participants with EEG procedures using a social story and practice net. As needed, evidence-based behavioral techniques were used to support compliance, including visual timers and token boards. FXS diagnosis was confirmed using Southern blot and PCR testing, identifying CGG triplet repeat expansions exceeding 200 repeats in the FMR1 gene (Liu et al., 2023).

**HBN**

All EEG data were collected using a 128-channel EEG recording system (Magstim EGI, Eugene, OR, USA). The EEG data were sampled at 500 Hz with a bandpass filter ranging from 0.1 to 100 Hz. The vertex (Cz) was used as the recording reference. Head circumference was measured for each participant to select an appropriately sized EEG net, and electrode impedances were maintained below 40 kΩ.

Diagnoses were derived using the Schedule for Affective Disorders and Schizophrenia Children's version (KSADS-COMP), a semi-structured DSM-5-based psychiatric interview administered by a licensed clinician. Clinician consensus diagnoses additionally incorporated supporting evidence from behavioral observations, family and intervention history, cognitive and academic testing, and parent, child, and teacher questionnaires (CBCL, SRS, SCQ, SCARED, among others). Diagnoses were assigned with levels of certainty: confirmed (full criteria met), presumptive or required confirmation (criteria likely met but limited by evaluation protocol), rule-out (insufficient evidence), or by history (previously diagnosed but not confirmed by HBN). For ASD specifically, the KSADS autism module served as the primary diagnostic component, with only a limited subset of participants additionally receiving gold-standard instruments such as the ADOS or ADI-R (Alexander et al., 2017).
HBN data are publicly available.

**MGF & Q1K**

The testing took place in a dark, soundproof experimental chamber in the Sainte-Justine university hospital center. Resting-state EEG was recorded using a high-density EEG system with 128 channels (Magstim EGI, Eugene, OR, USA). Signals were acquired by a G4 Macintosh computer using NetStation EEG Software (v. 4.5.4). Impedances were kept below 40 kΩ. Vertex (Cz) was used as online reference. EEG data was digitized at a sampling rate of 1000 Hz and analog bandpass-filtered from 0.1 to 500 Hz.
For all participants, the Autism Diagnostic Observation Schedule, Second Edition (ADOS-2) was performed unless it had been done in the past. Pathogenic CNVs were detected using the genome-wide chromosomal microarray analysis, which is routinely used in several units of the CHU Sainte-Justine. Sixty-nine recurrent and nonrecurrent pathogenic CNVs (50-500 Kb) were identified in this study sample (36 DEL and 33 DUP; 48% and 40% de novo respectively; 59 loci in total). The study protocol was reviewed and approved by the CHU Sainte-Justine Research Ethics Board. All methods were performed in accordance with the relevant guidelines and regulations. All the participants consented to participate in the study and signed the consent form. Eyes-open resting state was recorded for 4 min while participants were watching a movie without sound and subtitles to improve collaboration and reduce motion artifacts through gaze fixation on the screen.
MGF data are available from the corresponding author upon request.

**RDB**

Participants were diagnosed according to the DSM-5 criteria and were assessed using the Autism Diagnostic Interview - Revised (ADI-R), the Autism Diagnostic Observation Schedule (ADOS-2) and clinical reports from experts in the field, who made the final diagnostic decision.
The testing took place in an experimental chamber in the Robert Debré University Hospital Center, Paris, France. During the resting state, the subject remained seated and kept their eyes open for 2 minutes at the beginning and 1 minute at the end of the recording. The subject alternated between keeping their eyes open for 30 seconds (‘eyes-open’ condition) and closing their eyes for 30 seconds (‘eyes-closed’ condition). Additionally, the participant

blinked for 30 seconds ('blink' condition) and imitated chewing for 30 seconds ('chew' condition) to control the eye blink and jaw contraction artifacts.
HD-EEG was recorded using a 128-channel Geodesic EEG System 400 (GES 400) with NetStation EEG software and a 128-electrode HydroCel Geodesic Sensor Net (EGI, 2017). The sensor net was connected to a Net Amps 400 amplifier (EGI, 2017), which was set to '255 encoding mode' to record external Digital Input (DIN) events (triggers). The EEG data were amplified, low-pass filtered at 500 Hz, sampled at a frequency of 1000 Hz, and digitized.

**UNIGE**

Resting-state EEG recordings were acquired using 128-channel HydroCel Geodesic Sensor Nets and NetStation Acquisition EEG software (Electrical Geodesics System Inc., Eugene, OR, USA). The recordings were sampled at 1000 Hz with a band-pass filter ranging from 0 Hz to 100 Hz using the vertex as reference. Electrode impedances were maintained below 10 kΩ for the reference (REF) and common mode (COM) electrodes at the start of recordings.
Diagnosis of ASD was confirmed by a child psychiatrist (MS) according to the *Diagnostic and Statistical Manual of Mental Disorders* (5th ed., *DSM*-5) criteria (APA, 2013). To further confirm the diagnosis, all autistic participants underwent the Autism Diagnostic Observation Schedule (ADOS) and scored above diagnostic cut-off (Lord et al., 2000; Lord et al., 2012). Before inclusion in the cohort, TD children were also screened for any developmental concern, autistic features, somatic and psychiatric concerns, and for the absence of autism in first-degree family members. For more information about the cohort, see Latrèche et al. (2024).
Data access inquiries may be directed to the corresponding authors.

**SL**

Simons Searchlight EEG data used in this study are available to approved researchers via SFARI Base (dataset DS0000050: https://base.sfari.org/dataset/DS0000050). EEG/ERP data were collected during Phase 1 of the Simons Searchlight project at family meetings and at the lead site, Harvard University, using 128-channel HydroCel Geodesic nets by the Nelson Laboratory at Boston Children's Hospital. The dataset was released on November 14, 2018 (version 1, 13 GB). Access can be requested at https://base.sfari.org.
Genetic diagnoses were confirmed prior to enrollment through chromosomal microarray analysis (CMA), identifying deletions or duplications at the 16p11.2 locus. Participants were referred via medical genetics clinics, genetic counselors, and internet searches (Simons VIP Consortium, 2012). Where a proband was identified, cascade genetic testing was performed to identify other carrier family members.
